# Virus-Host Infection Dynamics of Marine Single-Celled Eukaryotes Resolved from Metatranscriptomics

**DOI:** 10.1101/093716

**Authors:** Mohammad Moniruzzaman, Louie L. Wurch, Harriet Alexander, Sonya T. Dyhrman, Christopher J. Gobler, Steven W. Wilhelm

## Abstract

Metatranscriptomics has emerged as a tool in microbial ecology that can resolve the functional landscape of both prokaryotes and eukaryotes within a community. In this study, we extend the potential of metatranscriptomics to probe active virus infections and virus-host relationships in marine systems. Polyadenylation-selected RNA-seq data were examined from microbial communities in two productive marine environments: a brown tide bloom event dominated by *Aureococcus anophagefferens* in Quantuck Bay, NY, and a diatom-dominated plankton community in Narragansett Bay, RI. Active infections by diverse giant viruses (NCLDVs) of algal and non-algal hosts were found at both sites. Ongoing infections of *A. anophagefferens* by a known *Mimiviridae* (AaV) were observed during both the peak and decline of the bloom. Bloom decline was also accompanied by increased activity for viruses other than AaV, including (+) ssRNA viruses. In Narragansett Bay, increased temporal reso’lution revealed active NCLDVs with both ‘boom-and-bust’ as well as ‘steady-state infection’-like ecologies. Statistical co-occurrence examinations of the dsDNA, ssRNA and dsRNA markers within the data revealed a broad spectrum of statistically strong and significant virus-host relationships that included both known as well as novel interactions. Our approach offers a method for screening the diversity and dynamics of active viral infections in natural systems and develops links between viruses and their potential hosts *in situ.*

**Significance:** Viruses are important partners in ecosystem scale ecology, yet their study to date is primarily limited to single virus-host infection models in the laboratory or limited to “potential-actions” derived from metagenomics analyses. Using metatranscriptomic sequences from polyadenylated-RNA selected samples, we have simultaneously captured information regarding eukaryotic diversity and active infection by viruses with dsDNA genomes, resulting in a statistical opportunity to predict “*who is infecting whom*”. This approach further provides concurrent insight regarding viruses with ssRNA and dsRNA genomes, capturing dynamics for the communities of viruses infecting single-celled eukaryotes. Given the central role of these plankton in global scale processes, our efforts result in a transformational step-forward regarding the study of *in situ* virus-host interactions.

## Introduction

Viruses that infect marine microbes are an integral component of aquatic ecosystems, with a diversity spectrum spanning the entire Baltimore classification scheme (1). The association of viruses with global-scale biogeochemistry, algal bloom termination events, and their impact on microbial community diversity has driven scientific research in virus ecology (2, 3). Amongst these predators, giant dsDNA viruses belonging to the Nucleocytoplasmic Large DNA Virus (NCLDV) group infect single-celled eukaryotes with diverse lifestyles (4) and are thought to be abundant in the world’s oceans (5). Individually some of these viruses have been shown to be potential drivers of algal bloom collapse (6, 7). However, only a few NCLDV-host-systems with established ecological relevance have been identified. As a specific example the environmental hosts of the *Mimiviridae,* isolated using *Acanthamoeba* in the laboratory, are yet to be confirmed (8).

Along with the NCLDVs, RNA viruses also comprise a major fraction of the marine viroplankton, infecting organisms ranging from diatoms and dinoflagellates to fish (9). However, little is known about the ecology and host range of RNA viruses; the first RNA virus infecting a marine single-celled eukaryote was only described in 2003 (10). In addition, recent evidence suggests that a large number of novel ssDNA virus families possibly infect yet to be characterized marine phytoplankton and zooplankton (11). Collectively, these observations illustrate the strong need to develop *in situ* approaches that link the marine virosphere to their hosts within the microbial eukaryotes.

The marine ecosystem consists of complex interactions among diverse organisms and their viruses. While studying individual host-virus systems remain critical to understand the molecular basis of interactions, studying the overall contribution of viruses in a dynamic network of organisms is hindered by methodological limitations. Culture independent approaches to study viruses, and especially viral communities, are challenging: “viromes” – large metagenomic datasets enriched with viral sequences, are usually generated by size exclusion (≤ 0.22 μm) of bacteria and small eukaryotes (2). This approach, however, largely removes the large virus particles that can range from 100 nM to ~1.5 μM (8). Moreover, by targeting DNA, these approaches examine only the presence of particles and not their activity. Additionally, RNA-containing virus particles must be targeted separately from DNA viruses, since common methods for virus enumeration (using dsDNA intercalating stains) and DNA-based metagenomics approaches cannot detect them (12). Consequently, there is a need for new toolsets to complement the current approaches and yet overcome the aforementioned issues to provide a more comprehensive picture of the viral dynamics

Here we examined metatranscriptomes from two highly productive marine sites on the east coast of USA – Quantuck Bay, NY and Narragansett Bay, RI. Quantuck Bay experiences recurring ecosystem disruptive brown tide blooms caused by the pelagophyte *Aureococcus anophagefferens* (13) which are shaped by a giant virus (AaV) (14, 15). Narragansett Bay is a highly productive system with seasonal diatom blooms, but a poorly described eukaryotic virus community. By employing selection for polyadenylation prior to sequencing, we were able to focus on active virus infections within eukaryotes. By using time-series data, we were able to capture emergent relationships of putative virus-host pairings and their ecological dynamics. This approach also allowed us characterize viruses actively infecting eukaryotes with diverse nucleic acid genomes (ssDNA, ssRNA and dsRNA).

## Results and discussion

### Temporal dynamics of active giant virus infections

To identify NCLDVs, we screened contig libraries generated at each study site for ten conserved NCLDV core genes (16). Reads from individual samples were mapped to the core gene contigs followed by library size normalization. At both sites, numerous contigs originating from NCLDV-specific Major Capsid Protein (MCP) were identified (Fig. 1). The abundance of reads mapped to MCP contigs was higher than the sum of mapped reads to all other NCLDV core gene contigs (Fig. 1) for all samples except QB-S3, confirming efforts suggesting MCP is a suitable marker for NCLDV diversity (Moniruzzaman et al. 2016) and that the MCP gene is highly expressed (17). Only distant homologs of MCP are present in *Poxviridae* (16) and there are no homologs in recently discovered Pandora- and Pithoviruses (8): to this end the ubiquity of this gene in all other NCLDV families makes it an excellent candidate for phylogenetic probing of metatranscriptomics data.

**Figure 1:**
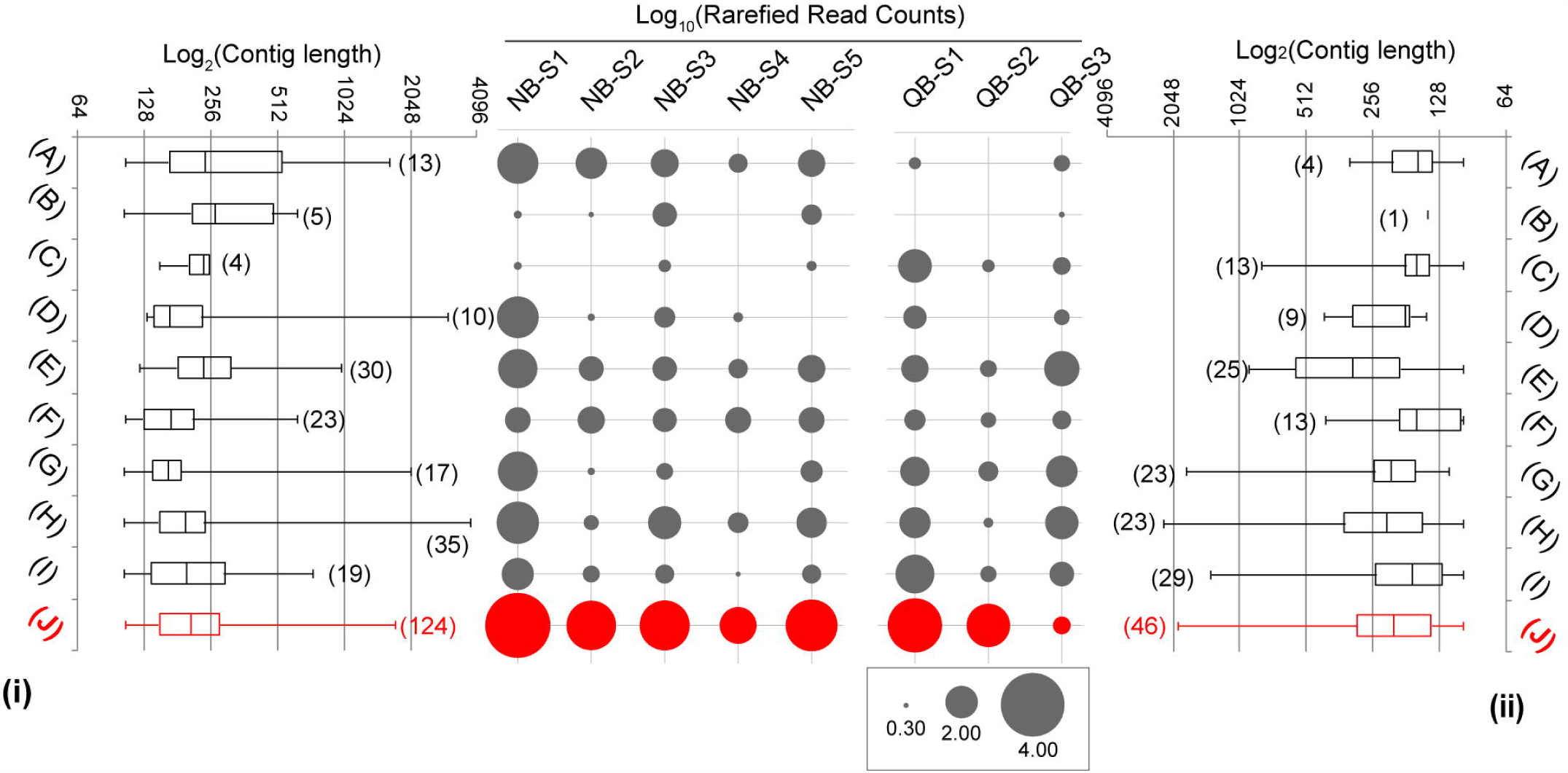
Abundance of 9 NCLDV core genes including major capsid protein (MCP) in terms of normalized read counts and number of contigs recovered (up to 100bp length) from Quantuck Bay (right panel) and Narragansett Bay (left panel). The box and whisker plots represent the range of the contig lengths with number of contigs recovered for each gene in brackets. The filled circles represent the rarefied abundances of each contig in each sample. No contigs could be detected from myristolyated envelope protein, a core NCLDV gene. Major capsid protein abundances are shown in red, while other core genes are presented in dark gray. Core genes are indicated on the X-axes as follows: A) A32 virion packaging ATPase, B) VLFT3 like transcription factor, C) Superfamily II helicase II, D) mRNA capping enzyme, E) D5 helicase/primase, F) Ribonucleotide reductase small subunit, G) RNA polymerase large subunit, H) RNA polymerase small subunit, I) B-family DNA polymerase, J) Major capsid protein.

We placed the MCP contigs on a reference phylogenetic tree and studied their relative expression levels in terms of a metric that we defined as ‘rarefied counts per kilobase’ (RCK) (details in Materials and methods). Phylogenetic placement of the contigs demonstrated that NCLDV members from *Mimiviridae* and *Phycodnaviridae* were consistently present in both Quantuck Bay and Narragansett Bay. At both sites the highest number of contigs fell within the *Mimiviridae* family, followed by *Phycodnaviridae* (Fig. 2). A large number of contigs had strong phylogenetic affinity to AaV as well as other alga-infecting members of the *Mimiviridae* clade. Their presence and relative abundance in these field surveys demonstrate the *Mimiviridae* are an important component of the marine virosphere and are as active as the better-studied *Phycodnaviridae* group.

**Figure 2:**
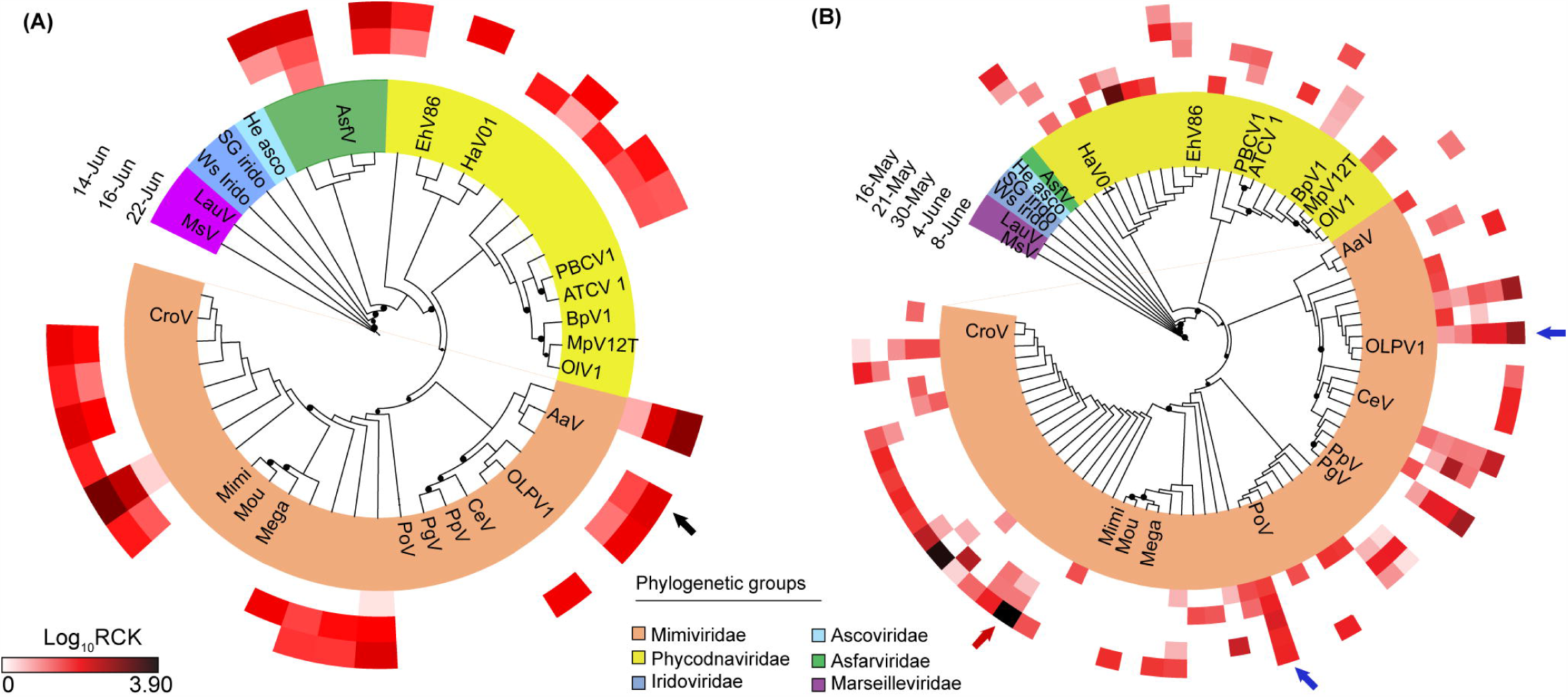
Phylogenetic placement of major capsid protein contigs from A) Quantuck Bay and B) Narragansett Bay on a reference tree of NCLDVs with icosahedral capsids. Node support (aLRT-SH statistic) >50% are shown as dark circles. Contigs upto 200bp are shown, with their expression level (rarefied read counts per kilobase – RCK) in individual samples as a heatmap on the outer rings. Notice that the placement trees contain both AaV reference sequence and the contig originating from AaV (marked with a black arrow). The reference sequences are shown in bold italic typeface. Abbreviations: MsV-Marseillevirs, LauV: Lausannevirus, Ws Irido: Weisenia iridescent virus, SG Irido: Singapore Grouper iridescent virus, He Asco: Heliothis virescens ascovirus, AsfV: Asfarvirus, EhV86: Emiliania huxleyi virus 86, HaV01: Heterosigma akashiwo virus 01, PBCV1: Paramacium bursaria Chlorella virus 1, ATCV 1: Acanthocystis turfacea Chlorella virus 1, BpV1: Bathycoccus parsinos virus 1, MpV12T: Micromonas pusilla virus 12T, OlV1: Ostreococcus lucimarinus virus 1, AaV: Aureococcus anophagefferens virus, CeV: Chrysochromulina ericina virus, PpV: Phaeocystis pouchetii virus, PgV: Phaeocystis globosa virus, PoV: Pyramimonas orientalis virus, Mega: Megavirus chilensis, Moumou: Moumouvirus goulette, Mimi: Mimivirus, CroV: Cafeteria roenbergensis virus.

Brown tide bloom samples collected on June 14 (QB-S1) and June 16 (QB-S2) represented the bloom peak with an *Aureococcus* count of ~2.28 x 10^6^ cells/mL and ~2.23 x 10^6^ cells/mL, respectively. The third sample, collected on June 22, represented the early stage of bloom demise, with an *Aureococcus* count of ~1.91 x 10^6^ cells/mL. We detected a persistent infection of *A. anophagefferens* by AaV across this sampling period. High stringency (similarity ≥ 97%) mapping of reads to the genome identified 1,368 and 604 reads that could be assigned to peak bloom samples QB-S1 and S2, respectively, after library size normalization, while 236 reads were mapped to the QB-S3 sample taken during bloom decline (Fig. S1). Across the entire genome, 15 AaV transcripts had more than 10 reads: roughly two thirds of these transcripts have no similarity to genes with currently known functions (ORFans) (Dataset S1). Highly expressed ORFans have also been recorded for Mimivirus: 17 of the top 20 most highly expressed genes were hypotheticals (17). These observations suggest these genes are active during infection by AaV and other NCLDVs, and represent important targets for future studies. Notably, the AaV MCP was among the most highly expressed functional genes, with 121 total reads mapped to this gene across the three *in situ* samples from Quantuck Bay. Both total reads mapped across the AaV genome (Fig. S1) and specifically to MCP gene (Fig. 2A) progressively declined throughout the sampling period, with the lowest number of reads mapped from S3. It may be that AaV activity was present, but reduced during the bloom decline stage — an observation that is supported by a recent study where AaV amplicons were only detected during the peak of the bloom (18). Overall these data reinforce the utility of MCP as a marker, since the MCP dynamics were consistent with data derived from the full AaV genomic analysis (Figure 2).

With five *in situ* samples over a period of approximately four weeks (Table S1), data from Narragansett Bay allowed us to observe the temporal dynamics of the NCLDVs. Some members from *Phycodna*- and *Mimiviridae* clades showed persistent evidence of infection over a prolonged period, while ‘boom-bust’ like relationships (4) were possibly present for other members (Fig. 2B). For example, a number of MCP contigs were consistently expressed (within an order magnitude) between samples across time points (*e.g*., blue arrows in Fig. 2B), an observation supporting the presence of infected hosts. While this scenario is consistent with a ‘slow-and-steady’ infection dynamic (19), it can also be explained by persistent infections of the plankton – where ongoing virus production does not necessarily lead to host (or at least total community) mortality (20). The expression of other phylogenetically distinct markers, however, reflected a ‘boom-and-bust’ like scenario (19), with the expression varying across several orders of magnitude between time points. One striking example of such putative ‘boom-and-bust’ scenario was a contig in the non-algal *Mimiviridae* family, where expression decreased by two orders of magnitudes from May 16 to May 21 and May 30 (Fig. 2B, red arrow).

### Viruses infecting single-celled eukaryotes beyond the NCLDVs

The marine virosphere is not limited to dsDNA viruses, as viruses containing all nucleic acid types (ss- and dsRNA as well as ssDNA) that infect marine single-celled eukaryotes have been described (9, 11). We extended our approach to detect the contigs that potentially originated from diverse RNA and DNA viruses other than NCLDVs. RNA viruses have a diverse size range, with Picornavirales particles as small as ~25-30 nm (21). Our sample collection method (Material and Methods) allowed detection of both ongoing virus infection (for DNA and RNA viruses) and cell-surface associated RNA viruses. It is important to mention that some (+) ssRNA viruses have poly-A tailed genomes (*e.g*., Picorna-and Togaviruses) even outside the host (22). Therefore, owing to their nature, the (+) ssRNA viral diversity captured by this approach might reflect both actively replicating and surface bound viruses.

Within our analyses, 579 and 599 contigs from Quantuck Bay and Narragansett Bay, respectively, were assigned to viruses other than NCLDVs. The majority of these contigs originated from (+) ssRNA viruses, with the main contributors coming from a yet unclassified group of viruses in the Picornavirales order (23). Unclassified Picornavirales contigs represented 62% of the total non-NCLDV viral contigs for Quantuck Bay and 74% of this group for Narragansett Bay. Marine Picornavirales have been shown to infect diatoms (*e.g., Chaetoceros* sp., *Asterionellopsis glacialis* and *Rhizosolenia setigera*) (9) and a marine fungoid protist, *Aurantiotrychium* (24). The closest phylogenetic relative of this group is *Marnaviridae,* which currently have only one member – HaRNAV, that infects the marine raphidophyte *Heterosigma akashiwo* (10). The second major group of (+) ssRNA viruses belonged to *Dicistroviridae* family, with 90 and 36 contigs from Quantuck Bay and Narragansett Bay, respectively (Fig. 3). Interestingly, the only dsRNA viruses detected in both locations were similar to viruses in the *Totiviridae, Partitiviridae* and *Hypoviridae* family, which are all known viruses of fungi (21). These viruses may be infecting fungi that are parasitic on algae, as have been proposed recently for samples collected in the Laurentian Great Lakes (25). While some ssDNA virus contigs from Quantuck Bay clustered with the *Nanoviridae* family, others from both locations did not form any definitive cluster with known circular DNA viruses, thus potentially representing previously undescribed circular DNA virus groups in the ocean (Fig. S2). No (−) ssRNA viral contigs were detected.

**Figure 3:**
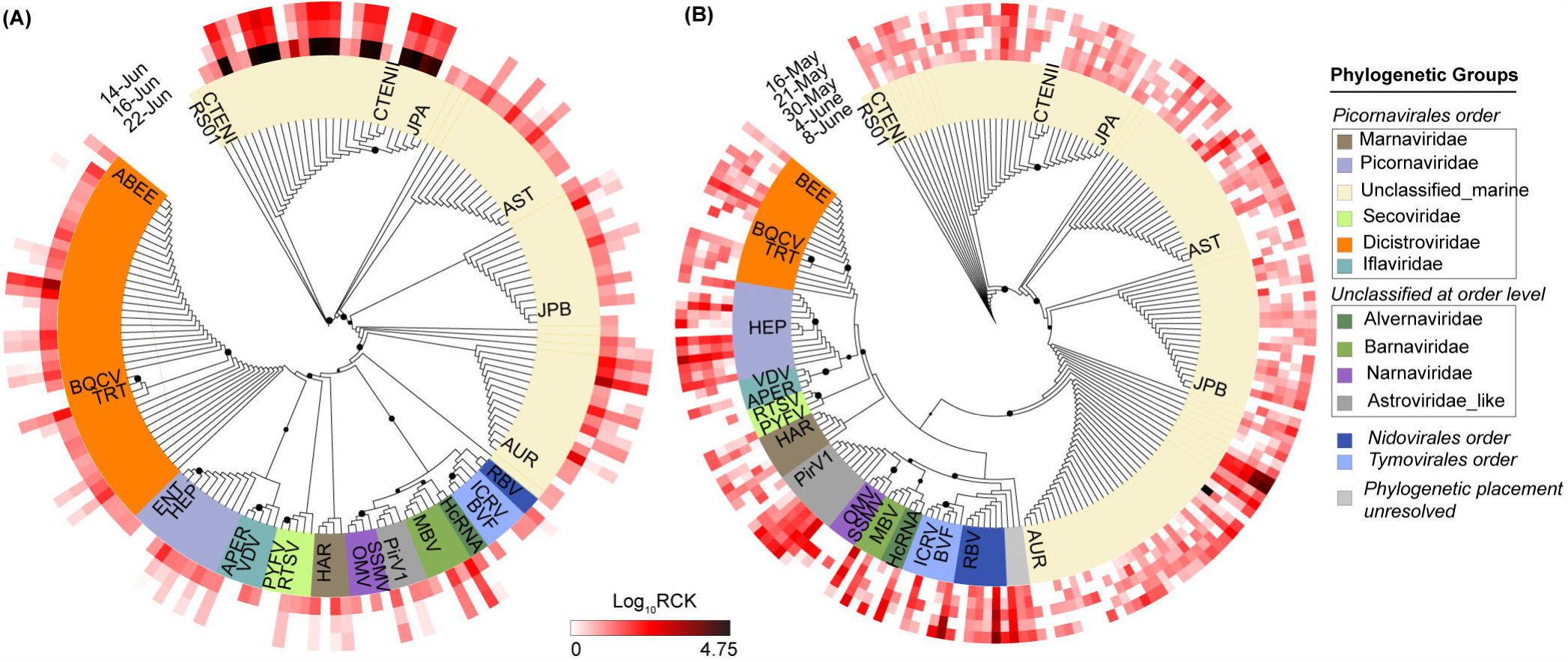
Phylogenetic placement of (+)ssRNA virus contigs harboring RNA dependent RNA polymerase (RdRP) motifs from A) Quantuck Bay and B) Narragansett Bay on reference trees. Node support (aLRT-SH statistic) >50% are shown as dark circles. Contigs up to 225bp are shown, with their expression level (rarefied read counts per kilobase – RCK) in individual samples as a heatmap on the outer rings. The reference sequences are shown in bold italic typeface. Complete name and other details of the reference sequences are presented in Supplementary Dataset 2.

To assess how the activity of virus groups changed over time, we measured the proportion of reads that mapped to different virus groups for each library. In Narragansett Bay, the majority of the virus reads originated from the unclassified marine Picornavirales and the *Dicistroviridae, Secoviridae* and *Picornaviridae* families across all the time points (Fig. S3). The unclassified marine Picornavirales group recruited from ~68% (NB-S1) to ~98% (NB-S5) of the non-NCLDV viral reads (Fig. S3). In Quantuck Bay, reads from both unclassified marine Picornavirales and ssDNA viruses dominated libraries during the first two time points (Fig. S4). However, a shift in the proportional abundance of virus reads was observed on the third day (QB-S3), when the unclassified marine Picornavirales became dominant (93% of the non-NCLDV virus transcripts, Fig. S4). Overall, 2.4% of the entire QB-S3 library (~4.3 million fragments) mapped to these unclassified Picornavirales, relative to 0.043% and 0.027% of reads for QB-S1 and QB-S2, respectively. This indicated a striking increase, concordant with the decline of the brown tide bloom. In parallel nutrient amended samples derived from water collected on the same date as QB-S3, the unclassified Picornavirales transcripts also increased by an order of magnitude relative to QB-S1 and QB-S2 (Data not shown) validating the increase in viral reads in QB-S3. Phylogenetic analysis confirmed these ssRNA viral contigs were consistent with the unclassified Picornavirales group (Fig. 3A). *Aureococcus* blooms are not mono-specific – they include diatoms, dinoflagellates and high densities (~10^4^ cells/ml) of heterotrophic protists (26). The striking increase in the unclassified Picornavirales could be related to infection of a host that co-occurs with *Aureococcus* and the potential shift in competition that might occur during *Aureococcus* bloom decline. These observations suggest a broad ecological role for viral infection during phytoplankton bloom decline, which would not have been resolved with targeted studies of AaV or metagenomic approaches. Taken together, the apparent dynamics and abundance of this unclassified Picornavirales suggest this group is a major component of the marine virioplankton, and strengthens previous observations that the Picornavirales phylogenetically distinct from the established families can be dominant members in different marine environments (12, 27). Owing to their small size, detection and quantification of RNA viruses and ssDNA viruses pose significant technical challenges (27). Our results, however, clearly point to the power of screening metatranscriptomes for the simultaneous analysis of the dynamics of a large cross-section DNA and RNA viruses.

Eighteen of the assembled contigs (9 from each site) were >7,000 bp and had best hits to different Picornavirales members. Phylogenetic analysis based on the RNA-dependent RNA polymerase (RdRp) gene and feature analysis of existing (+) ssRNA virus genomes suggested that these contigs are complete or near-complete Picornavirales genomes (Fig. 4). Sixteen of these genomes were dicistronic – they harbored two ORFs coding for structural and non-structural proteins, while the remaining 2 were monocistronic (Fig. 4), revealing differences in genome architecture among group members (Fig. 4). Remarkably, one virus (N_001) had a reverse orientation of the genes with the first ORF encoding for the structural protein, which is unusual for dicistronic Picornavirales (28). In addition, a glycosyl transferase domain was found in N_137 (Fig. 4). To our knowledge, the presence of glycosyltransferase domains has only been reported in members of the *Endornaviridae* family dsRNA viruses (29).

**Figure 4:**
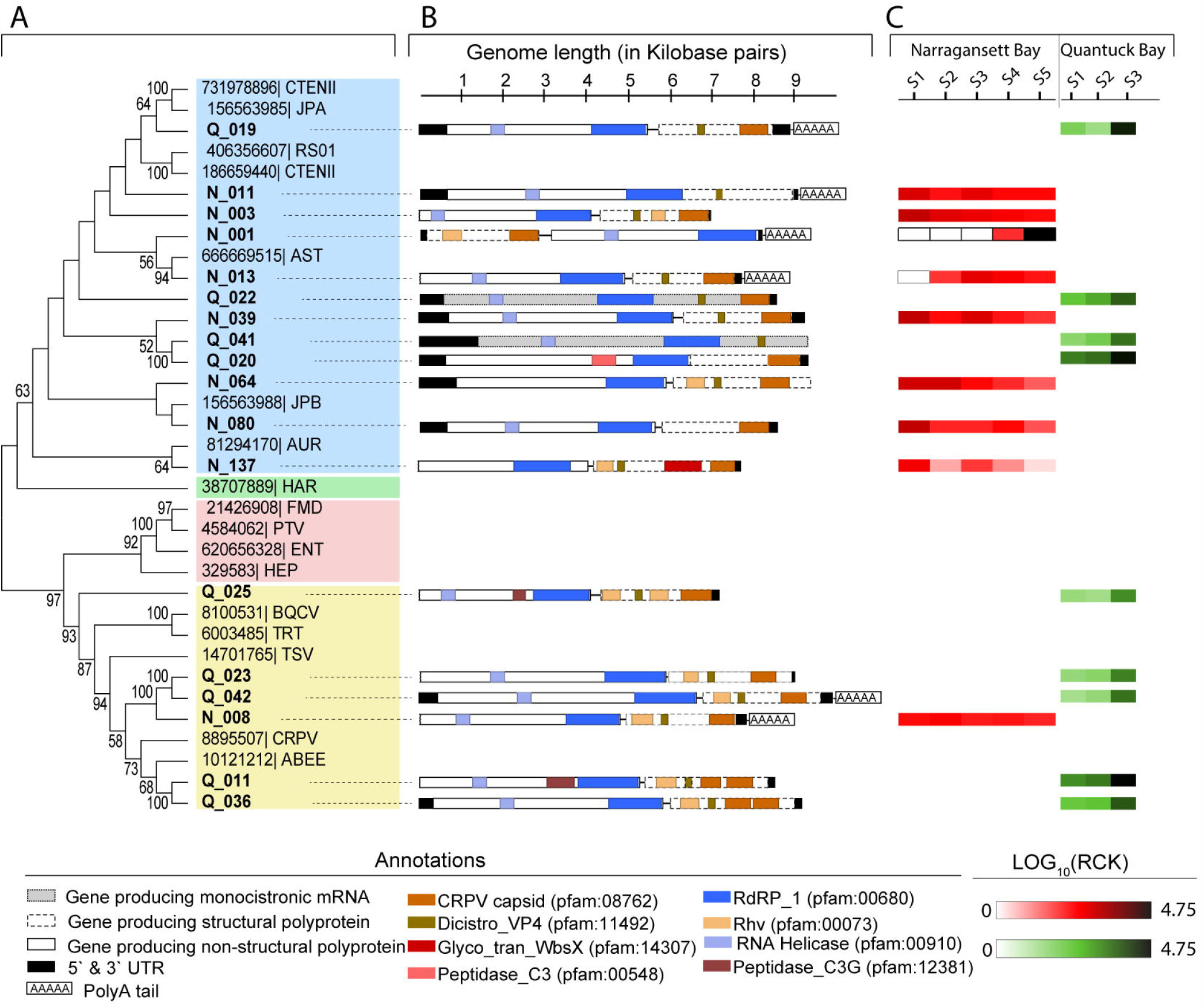
Complete or near-complete Picornavirales genomes recovered from both Quantuck Bay and Narragansett Bay study sites. Panel (A) shows the phylogenetic classification of these contigs in a topology-only maximum likelihood tree, with contigs from Quantuck Bay having prefix ‘Q_’ and contigs from Narragansett Bay having prefix ‘N_’. Panel (B) shows the genome architecture of these contigs with protein domains and putative CDSs. Panel (C) shows the expression level of these (rarefied read counts per kilobase – RCK) viruses in across different *in situ* samples.

We also tracked the dynamics of these *de novo* assembled genomes by mapping the data collected over spatiotemporal gradients. All the (+) ssRNA virus genomes from Quantuck Bay samples showed higher relative expression during bloom decline (QB-S3) compared to the time points corresponding to the bloom peak (Fig. 4, panel C). N_001, a candidate virus from Narragansett Bay, was not present in the first three time points. Its expression was only observed during the fourth sampling point, which was followed by a dramatic increase during the last sampling point, when it recruited ~0.55% of the reads from the entire library (Fig. 4, panel C). The closest known phylogenetic relative of N_ 001 is a virus infecting diatom *Asterionellopsis glacialis* (Fig. 4), suggesting the putative host may be a diatom. Narragansett Bay was experiencing a spring diatom bloom during the sampling period with “boom—bust” abundance cycles in the relative abundance of putative diatom hosts (30), consistent with these observed viral dynamics.

### Who is infecting whom? Resolving virus-host relationships using metatranscriptomics

This study presented the opportunity to evaluate potential relationships among diverse single-celled eukaryotes and their pathogens, with the established *AaV-Aureococcus* association acting as a *de facto* internal standard. Transcripts from DNA viruses must originate within the host cells, and thereby for a particular host-virus pair, a significant and strong positive correlation is to be expected for gene expression. Building on this idea, host gene expression of at least a subset of the host’s genome is a prerequisite to observe gene expression of a virus specific to that host, as evidenced by transcriptomic landscape of host-virus dynamics in culture (17, 31) and induced blooms in mesocosms (32). To expand our data, we also took advantage of concurrent nutrient amendment studies in mesocosms (see Methods and Materials) which provided additional samples for our analyses.

We inspected statistical co-occurrences among the contigs containing virus and eukaryote-specific marker genes based on their expression patterns. Since the polyadenylated-selected metatranscriptomes are largely depleted of ribosomal RNA marker genes, we employed functional genes suitable for phylogenetic analysis. Expression of MCP (dsDNA NCLDVs), RNA dependent RNA polymerase (RdRP) (RNA viruses) and viral replicase (ssDNA viruses) were compared to the functional eukaryotic marker gene RNA polymerase II large subunit (RPB1, Fig. S5), a candidate gene to resolve the phylogenetic history of different eukaryotic lineages (33, 34). Hierarchical clustering of a Pearson’s correlation matrix followed by SIMPROF analysis (35) was used to detect statistically distinct clusters which contained both virus and eukaryotic marker genes that could be classified into phylogenetic groups. This analysis produced several statistically distinct clusters harboring both viral and eukaryotic contigs (Fig. 5). A single cluster (Fig. 5A(ii)) harbored both AaV and *Aureococcus,* validating that established ecologically relevant relationships between viruses and their hosts can be retrieved using this approach.

**Figure 5:**
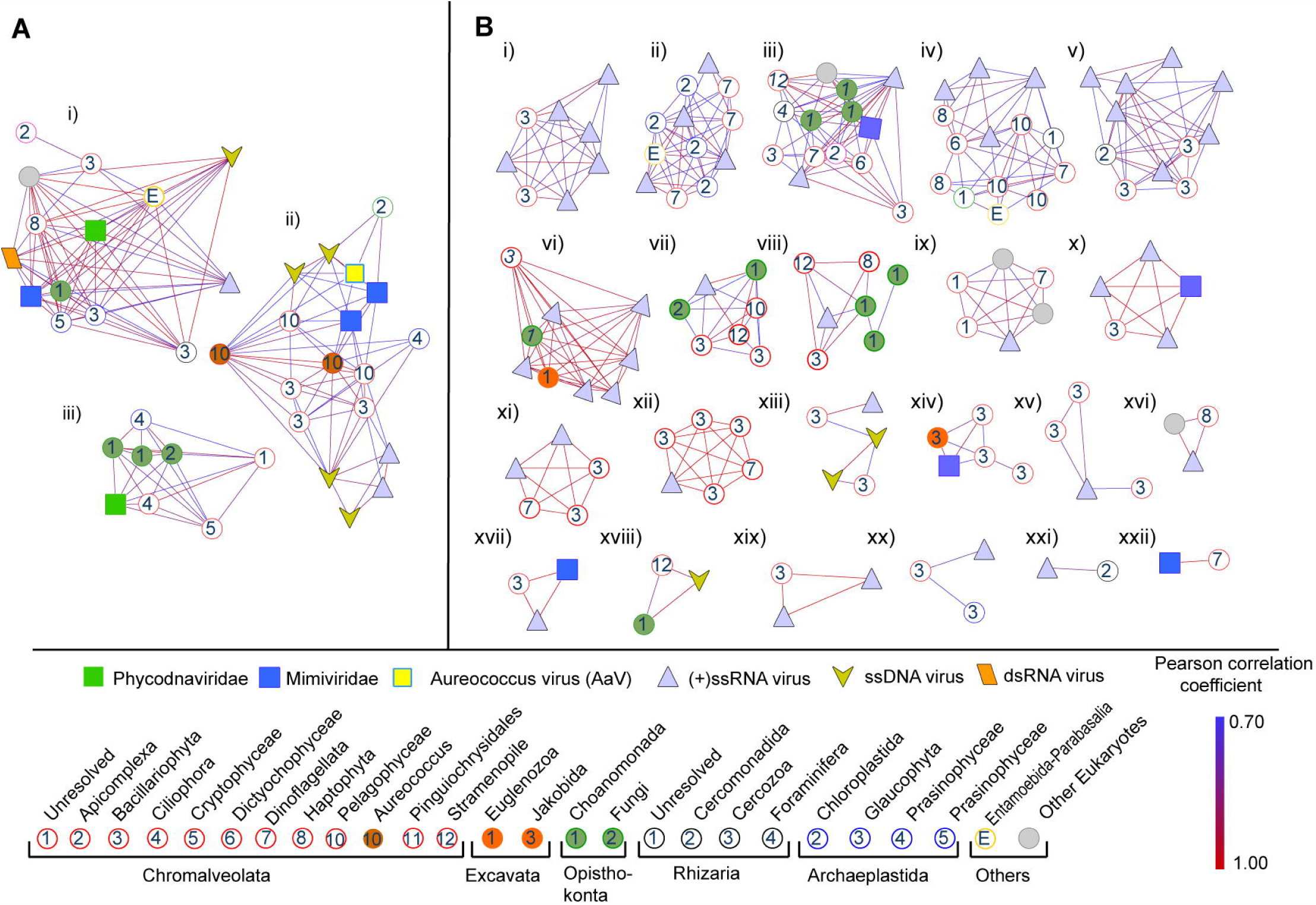
Representative SIMPROF clusters containing both viral and eukaryotic members from A) Quantuck Bay and B) Narragansett Bay. Contigs are shown as nodes and the correlations as the connecting edges. Phylogenetic classifications of the contigs are shown in the bottom panel. *Aureococcus anophagefferens* (dark brown circles) and Aureococcus anophagefferens virus (AaV) (bright yellow square) are in cluster A(ii).

Close inspection revealed other interesting relationships among the coexisting eukaryotic and viral components. Cluster A(ii), while containing both *Aureococcus* and AaV, also contained another *Mimiviridiae* member, several ssDNA and (+) ssRNA viral contigs along with eukaryotes belonging to prasinophyceae and pelagophyceae (Fig. 5). The possibility of *Aureococcus* being infected by more than one virus type cannot be discounted (and indeed is perhaps likely). Moreover, the potential for AaV to infect closely related pelagophytes remains a possibility (although this has not been seen in lab studies) (36). One cluster, A(i), which contained both a *Phycodna*- and a *Mimiviridae* member, also included a RPB1 contig phylogenetically placed in the Cercozoa group (Fig. 5). Although no cercozoan host-NCLDV pairs currently exist in culture, a recent study showed integration of NCLDV genes in the genome of a cercozoan *Bigelowella natans* (37). This integrated NCLDV in the *B. natans* genome potentially belongs to the *Phycodnaviridae*, as revealed by phylogenetic analysis of the MCP gene.

Similar clusters of phylogenetically distinct eukaryotes and viruses were also found in Narragansett Bay. For example, cluster B(iii) contained a *Mimiviridae* and several ssRNA viral contigs connected to choanomonada, stramenopile, diatom and dinoflagellate members (Fig. 5). The majority of the eukaryotic contigs belonged to diatoms and dinoflagellates in the Narragansett Bay samples, which reflects the composition of single-celled eukaryotes in this site (30). A large number of contigs having phylogenetic affinity to choanomonada were found in both Quantuck Bay and Narragansett Bay locations and were represented in several of the representative SIMPROF clusters (Fig. 5). While larger networks of viruses and eukaryotes also existed, clusters with fewer members revealed more specific relationships. For example, cluster B(xiv) contained one *Mimiviridae,* one jakobida (heterotrophic flagellate) and several diatom contigs (Fig. 5). To date the obligate heterotrophs known to be infected by *Mimiviridae* members are *Cafeteria roenbergensis* (38), *Acanthamoeba* (39) and *Vermaamoeba* spp (40). Additionally; Cluster B(xviii) harbored a ssDNA virus, a stramenopile and a choanomonada member, while cluster B(xxii) revealed a one-to-one relationship between a *Mimiviridae* and a dinoflagellate (Fig. 5). Only one dinoflagellate – *Heterocapsa circularisquama,* has been shown to be infected by a NCLDV (41), so this potential host-virus pair is of particular note. Cluster B(x) and B(xvii) consisted of *Mimiviridae,* diatoms and ssRNA viruses. No diatom is yet known to be infected by a NCLDV, although a large number of ssRNA viral contigs in our study are phylogenetically close to diatom-infecting RNA viruses in the unclassified marine picornavirales group (Fig. 3). A number of clusters (*e.g*., B(xii)) were enriched with both ssRNA virus and diatom contigs. These relationships between ssRNA viruses and the eukaryotes needs to be interpreted with caution, however, as these contigs might originate both from free virus particles and/or viruses within hosts.

Several clusters also contained fungal contigs along with other eukaryotes – pointing to the possibility of broad parasitic relationships with phytoplankton and other single-celled eukaryotes. The AaV-*Aureococcus* cluster A(ii) harbored a fungal contig and a *Barnaviridae* member – a virus family with fungi as the only known hosts (Fig. 5) (21). Several other clusters, *e.g*., A(iii) and B(vii) also contained fungal contigs. While such observations are not definitive, they point to the existence of parasitic relationships resulting in complicated ecological interactions involving unicellular eukaryotes, fungi and fungal viruses in marine ecosystems (Edgar *et al*., 2016).

Increased sample resolution in the future will resolve more statistically robust relationships, which can further narrow potential interacting partners. One limitation of reference independent assembly of high throughput data is fragmented contigs originating from same transcript – which is illustrated by two *Aureococcus* specific RPB1 contigs in cluster A(ii) that originated from a single coding sequence. Increased sequencing, along with the continued development of assembly tools will provide better resolution to these relationships. These limitations notwithstanding, the analysis provides a ‘proof-of-principle’ for inferring the complex relationships among diverse unicellular eukaryotes and their viruses using metatranscriptome data.

## Conclusion

In this study we demonstrate how metatranscriptomics can provide a unique view of the marine virosphere by simultaneously detecting multiple viral infections across the landscape of the eukaryotic plankton within an ecosystem setting. This effort can largely overcome previous technical limitations involved in the study of different viral groups, owing to their size range and genome type, within the same sample. In the last two decades we have learned much regarding the diversity and dynamics of the phages in the ocean, but the eukaryotic virosphere has remained elusive, with little known about who is infecting whom in the environment (42). As demonstrated in our study, analyzing the vast wealth of information captured by metatranscriptomics, in a statistical framework, can be an important step towards answering this vital question.

## Acknowledgements

The authors thank Gary LeCleir and Eric Gann for assistance in data collection and analyses. This work was funded by grants from the Gordon and Betty Moore Foundation (Grant EMS4971 to SWW), The National Science Foundation (OCE-1061352) and the *Kenneth & Blaire Mossman Endowment* to the University of Tennessee (SWW). Additional funding was provided by the NOAA ECOHAB Program (NA15NOS4780199 to STD and CJG).

## Materials and Methods

### Experimental design

#### Quantuck Bay

Samples were collected from a brown tide bloom in Quantuck Bay (Latitude = 40.806395; Longitude = −72.621002), NY that occurred from late May to early July, 2011, covering the initiation, peak and demise of the bloom. Samples collected on June 14 (BT-S1) and June 16 (BT-S2) represented the peak of the bloom, while sample collected on June 22 (BT-S3) represented the initial phase of bloom decline. *Aureococcus* cells were counted from glutaraldehyde (1% final v/v) fixed whole water samples using an enzyme-linked immunosorbent assay (ELISA) with a monoclonal antibody as described previously (43). *In situ* samples from June 22^nd^ (3rd sampling point) was also used to carry out nutrient amendment experiments. Briefly, bottles were filled with natural sea water from the bloom and were amended with 25 μM ammonium only (+N), 4 μM phosphate only (+P), and 25 μM ammonium and 4 μM phosphate (+N&P) in triplicate. Three additional bottles with no nutrient addition were used as control. The samples were then incubated for 24 hours in a floating chamber at 0.5 m in eastern Shinnecock Bay at the Stony Brook — Southampton Marine Science Center under one layer of neutral density cover to mimic the light and temperature levels of Quantuck Bay. Samples for *Aureococcus* cell density measurement and total RNA extraction were collected at T=0 and T=24 hours. Approximately 25 ml of natural seawater from each of the *in situ* and nutrient amendment samples were pre-filtered through 5 μm polycarbonate (PC) filters and collected on 0.2 μm PC filters. The samples were flash frozen immediately after filtration and transferred to −80°C. Prior RNA extraction, CTAB buffer (Teknova, CA, USA) amended by polyvinylpyrrolidone (1% mass/vol) was added to each of the samples.

#### Narragansett Bay

The details sampling procedure is described in Alexander et al (30). Briefly, samples were collected from a long term sampling site in Narragansett Bay (41°34′12″ N, 71°23′24″ W) during 2012 on May 16 (NB-S1), May 21 (NB-S2), May 30 (NB-S3), June 4 (NB-S4) and June 8 (NB-S5). Sample collection and processing was completed within 0830 and 0900 local time to reduce the influence of diel signals. 2.0 L of water from each sample was filtered on 5.0-μm pore size PC filters using a peristaltic pump. The filters were snap frozen in liquid nitrogen and stored at −80°C until RNA extraction. Water collected along with NB-S3 was also used for nutrient amendment experiments. For this, triplicate 2.5L bottles were filled with water pre-filtered through a 200-μm mesh and amended with specific nutrients to create +N, +P, −N, −P treatments alongside an ambient control. The +N and +P treatments were designed to eliminate nitrogen and phosphate stress signals, whereas the −N and −P treatments were designed to drive the treatments towards each limitation respectively by skewing the nutrient ratios (Alexander *et al*., 2015). N and P amendment concentrations were ~10-fold the seasonal average N and P concentrations measured at the station II in the surface waters of Narragansett Bay. The +P and +N amendment contained 3 μM phosphate and 10 μM nitrate, respectively. The −P amendment contained 10 μM nitrate, 68 μM silica, 4.6 μM iron and f/5 vitamins. The −N treatment was amended with 3 μM phosphate, 68 μM silica, 4.6 μM iron and f/5 vitamins. The f/5 media ratios (44) were followed for silica, iron and vitamin amendments. Bottles were incubated for 48 hours in a flow-through incubator at ambient temperature and photosynthetically active radiation. After the end of the incubation, treated and control samples were filtered and stored for RNA extraction in the same manner for the *in situ* samples.

### RNA extraction and sequencing

Quantuck Bay samples were extracted using the UltraClean^®^ Plant RNA Isolation Kit (MO BIO Laboratories, CA, USA) according to manufacturer’s protocol. RNA samples were quantified spectrophotometrically and were sequenced in the Columbia Sequencing Center (NY, USA) using Illumina™ HiSeq™ platform with poly-A enrichment at a depth of ~50 million 100bp single end reads. Two more replicate samples were sequenced from June 22 (QB-S3) at a depth of 100 million reads. For the present study, these biological replicates from QB-S3 were pooled together prior to further analysis. The field sequence data reported in this paper have been deposited in the National Center for Biotechnology Information Sequence Read Archive, *www.ncbi.nlm.nih.gov/sra* (Narraganset Bay accession no. SRP055134; Quantuck Bay accession no. SRP072764).

For Narragansett Bay, replicate filters from each treatment and *in situ* samples were pooled, representing 6 L of water for each sample. RNA was extracted using RNeasy Mini Kit (Qiagen, Hilden, Germany) according to a modified yeast RNA extraction protocol. Briefly, lysis buffer and RNA-clean zircon beads were added to the filter. Samples were then vortexed for 1 min, placed on ice for 30 s, and then vortexed again for 1 min. The resulting RNA was eluted in water and possible DNA contamination was removed using a TURBO DNA-free Kit (Thermo Fisher Scientific, MA, USA). RNA from each triplicate was pooled by sample or treatment. >1,000 ng RNA from each sample then went through a poly-A selection using oligo-dT beads followed by library preparation with TruSeq RNA Prep Kit (Illumina, CA, USA). The samples were sequenced with an Illumina™ HiSeq2000™ at the Columbia University Genome Center to produce ~60 million; 100bp paired-end reads per sample.

### Read assembly and screening for virus and eukaryote specific contigs

Sequence reads from both locations were quality trimmed (stringent trimming (quality score ≤0.03), No ‘N’s allowed, 70bp size cutoff) in CLC genomics workbench 8.0 (Qiagen, Hilden, Germany). The data from all the Quantuck Bay samples were combined and assembled together, and a similar assembly procedure was also followed for the Narragansett Bay specific samples. This resulted in 2,455,926 contigs for Quantuck Bay and 9,525,233 contigs for Narragansett Bay at a 100bp size cut-off.

For selecting contigs specific to Major Capsid proteins of NCLDV, a HMM profile was created after aligning the MCP sequences from complete giant virus genomes and several reported MCP genes available in NCBI. The HMM profile was queried against the translated contig libraries to select the putative MCP candidate contigs using HMMER (45). For selecting eukaryotic RPB1 contigs, HMM profile specific to domain “RPB1-C-term (NCBI CDD ID: cd02584)” and “RPB1-N-term (NCBI CDD ID: cd02733)” was used to query the contig libraries. All the MCP and RPB1 candidate contigs detected in this manner were queried against NCBI Refseq database and only contigs with first BLASTx hits (e-value cut-off ≤10^−3^) to MCP and RPB1 were kept for further analysis.

To detect contigs originating from viruses other than NCLDVs, we combined the proteins derived from all the viruses having algal, fungal and protozan hosts available on NCBI database. This protein database was queried against the contig libraries using tBLASTn with an e-value cut-off of ≤10^−3^. All the candidate contigs screened by this procedure were then queried against NCBI Refseq database using BLASTx. Only contigs having topmost hits to different viruses were kept for further analysis. All these contigs had best hits to diverse eukaryotic viruses- no hits to prokaryotic viruses were detected. This is probably due to the fact that the samples were poly-A selected prior to sequencing. These virus contigs were binned into distinct viral groups according to their best BLASTx hits. Percentage of reads recruited to individual viral groups was calculated for determining proportional abundance of different viral groups over different time points. For detailed phylogenetic analysis of ssRNA and ssDNA viruses, subset of these contigs harboring RdRP (pfam id: PF05183) and viral replicase (pfam: PF03090) motif were selected using HMM profile specific to these motifs.

### Genomic and phylogenetic analysis

Reference sequences for MCP (giant viruses), RdRP (RNA viruses), viral replicase (ssDNA virus) and RPB1 (eukaryotes) were downloaded from NCBI Refseq database. A number of RPB1 sequences representing several eukaryotic groups were also collected from Marine Microbial Eukaryotes Transcriptome Sequencing Project (MMETSP) (46) peptide collections, for which no representative genomes are available. The reference sequences were aligned in MEGA 6.0 (47). Maximum likelihood phylogenetic trees were constructed in PhyML (48) with LG model, gamma shape parameter and frequency type estimated from the data. aLRT SH-like statistic was calculated for branch support. The eukaryotic classification scheme by Adl *et al*. (49) was followed. Selected contigs were translated to amino acid sequences and were placed on the reference trees in a maximum likelihood framework using pplacer (50). The placement files were converted to trees with pendant edges showing the best placement of the contigs using ‘guppy’ tool of pplacer. The placement trees were visualized and annotated using iTOL interface (51).

ORFs were predicted on the complete or near-complete Picornavirales genomes using CLC genomic workbench 8.0 (www.clcbio.com). The genome annotation with predicted features was assisted by pfam (52) and Conserved Domain Database (CDD) (53) search. The genomes are submitted in the NCBI database under the accession numbers: KY286099 — KY286107 and KY130489 — KY130497.

### Statistical analysis

Quality trimmed reads were mapped to the selected viral and eukaryotic contigs from individual read libraries with high stringency (97% identify, 70% length fraction matching) in CLC genomics workbench 8.0. The read mapping values were normalized by library size and length and expressed as rarefied counts per kilobase (RCK). RCK values of viral and eukaryotic contigs >225 base pairs were converted into matrices separately for Quantuck Bay and Narragansett Bay datasets, which included mapping statistics from both *in situ* and nutrient amendment libraries. Group averaged hierarchical clustering was performed on these matrices using Pearson’s correlation coefficient in PRIMER 7.0. SIMPROF test (35) was applied on the clusters with 5% significance level and 1000 permutations to identify statistically distinct clusters. Selected clusters were visualized and annotated in Cytoscape 3.0 (54).

## References

1. Breitbart M (2012) Marine viruses: truth or dare. Ann Rev Mar Sci 4:425–448.

2. Brum JR, et al. (2015) Patterns and ecological drivers of ocean viral communities. Science 348(6237).

3. Weitz JS & Wilhelm SW (2012) Ocean viruses and their effects on microbial communities and biogeochemical cycles. F1000 Biol Rep 4:17.

4. Short SM (2012) The ecology of viruses that infect eukaryotic algae. Environ Microbiol 14(9):2253–2271.

5. Ogata H, Monier A, & Claverie J-M (2011) Distribution of Giant Viruses in Marine Environments. Global Change: Mankind-Marine Environment Interactions, eds Ceccaldi H.J, Dekeyser I, Girault M, & Stora G (Springer Netherlands), pp 157–162.

6. Lehahn Y, et al. (2014) Decoupling physical from biological processes to assess the impact of viruses on a mesoscale algal bloom. Curr Biol 24(17): 2041–2046

7. Gastrich M, et al. (2004) Viruses as potential regulators of regional brown tide blooms caused by the alga,Aureococcus anophagefferens. Estuaries 27(1):112–119.

8. Abergel C, Legendre M, & Claverie J-M (2015) The rapidly expanding universe of giant viruses: Mimivirus, Pandoravirus, Pithovirus and Mollivirus. FEMS Microbiol Rev 39(6):779–796

9. Lang AS, Rise ML, Culley AI, & Steward GF (2009) RNA viruses in the sea. FEMS Microbiol Rev 39 33(2):295–323.

10. Tai V, et al. (2003) Characterization of HaRNAV, a single-stranded RNA virus causing lysis of Heterosogma akashiwo (Raphidophyceae). J Phycol 39(2):343–352.

11. Labonte JM & Suttle CA (2013) Previously unknown and highly divergent ssDNA viruses populate the oceans. ISME J 7(11):2169–2177.

12. Steward GF, et al. (2013) Are we missing half of the viruses in the ocean? ISME J 7(3):672–679.

13. Gobler CJ, et al. (2011) Niche of harmful alga Aureococcus anophagefferens revealed through ecogenomics. Proc Natl Acad Sci USA. 108(11):4352–4357

14. Gastrich MD, Anderson OR, Benmayor SS, & Cosper EM (1998) Ultrastructural analysis of viral infection in the brown-tide alga, Aureococcus anophagefferens (Pelagophyceae). Phycologia 37(4):300–306.

15. Gastrich MD, Leigh-Bell JA, Gobler C, Anderson OR, & Wilhelm SW (2004) Viruses as potential regulators of regional brown tide blooms caused by the alga, Aureococcus anophagefferens: a comparison of bloom years 1999-2000 and 2002. Estuaries 27(1): 112–119.

16. Yutin N, Wolf Y, Raoult D, & Koonin E (2009) Eukaryotic large nucleo-cytoplasmic DNA viruses: Clusters of orthologous genes and reconstruction of viral genome evolution. Virol J 6(1):223.

17. Legendre M, et al. (2010) mRNA deep sequencing reveals 75 new genes and a complex transcriptional landscape in Mimivirus. Genome Res 20(5):664–674.

18. Moniruzzaman M, et al. (2016) Diversity and dynamics of algal Megaviridae members during a harmful brown tide caused by the pelagophyte, Aureococcus anophagefferens. FEMS Microbiol Ecol. 92: fiw058

19. Brussaard CPD (2004) Viral control of phytoplankton populations—a review. J Euk Microbiol 51(2):125–138.

20. Floge SA (2014) Virus infections of the eukaryotic marine microbes. PhD Dissertation (The University of Maine, Maine).

21. Wilson WH, Van Etten, J.L., Schroeder, D.C., Nagasaki, K., Brussaard, C., Bratbak, G., Suttle, C. (2012) Virus Taxonomy: Ninth Report of the International Committee on Taxonomy of Viruses. ed Andrew M.Q. King EL, Michael J. Adams, Eric B. Carsten (Elsevier London, UK), pp 10–20.

22. Shatkin AJ (1974) Animal RNA viruses: genome structure and function. Ann Rev Biochem 43(1):643–665.

23. Culley AI, Lang AS, & Suttle CA (2003) High diversity of unknown picorna-like viruses in the sea. Nature 424(6952):1054–1057.

24. Takao Y, Mise K, Nagasaki K, Okuno T, & Honda D (2006) Complete nucleotide sequence and genome organization of a single-stranded RNA virus infecting the marine fungoid protist Schizochytrium sp. J Gen Virol 87(Pt 3):723–733.

25. Edgar RE, et al. (2016) Adaptations to photoautotrophy associated with seasonal ice cover in a large lake revealed by metatranscriptome analysis of a diatom bloom. J Gt Lakes Res 42 (5):1007–1015.

26. Sieracki ME, et al. (2004) Pico- and nanoplankton dynamics during bloom initiation of Aureococcus in a Long Island, NY bay. Harmful Algae 3(4):459–470.

27. Miranda JA, Culley AI, Schvarcz CR, & Steward GF (2016) RNA viruses as major contributors to Antarctic virioplankton. Environ Microbiol. 18(11):3714–3727.

28. Greninger AL & DeRisi JL (2015) Draft Genome Sequence of Laverivirus UC1, a Dicistrovirus-like RNA virus featuring an unusual genome organization. Genome Announc 3(4):e00656–00615.

29. Song D, Cho WK, Park S-H, Jo Y, & Kim K-H (2013) Evolution of and horizontal gene transfer in the Endornavirus genus. PLoS ONE 8(5):e64270.

30. Alexander H, Jenkins BD, Rynearson TA, & Dyhrman ST (2015) Metatranscriptome analyses indicate resource partitioning between diatoms in the field. Proc Natl Acad Sci USA 112(17):E2182–E2190.

31. Rowe JM, et al. (2014) Global Analysis of Chlorella variabilis NC64A mRNA profiles during the early phase of Paramecium bursaria Chlorella Virus-1 infection. PLoS ONE 9(3):e90988.

32. Pagarete A, et al. (2011) Unveiling the transcriptional features associated with coccolithovirus infection of natural Emiliania huxleyi blooms. FEMS Microbiol Ecol 78(3):555–564.

33. Stiller JW & Hall BD (1997) The origin of red algae: implications for plastid□evolution. Proc Natl Acad Sci USA 94(9):4520–4525.

34. Malik SB, et al. (2011) Phylogeny of parasitic parabasalia and free-living relatives inferred from conventional markers vs. Rpb1, a single-copy gene. PLoS One 6(6):e20774.

35. Clarke KR, Somerfield PJ, & Gorley RN (2008) Testing of null hypotheses in exploratory community analyses: similarity profiles and biota-environment linkage. J Exp Mar Biol Ecol 366(1-2):56–69.

36. Gobler CJ, Anderson OR, Gastrich MD, & Wilhelm SW (2007) Ecological aspects of viral infection and lysis in the harmful brown tide alga Aureococcus anophagefferens. Aquat Microb Ecol 47(1):25–36.

37. Blanc G, Gallot-Lavallee L, & Maumus F (2015) Provirophages in the Bigelowiella genome bear testimony to past encounters with giant viruses. Proc Natl Acad Sci U S A 112(38):E5318–5326.

38. Fischer MG, Allen MJ, Wilson WH, & Suttle CA (2010) Giant virus with a remarkable complement of genes infects marine zooplankton. Proc Natl Acad Sci USA 107(45):19508–19513.

39. Abrahão JS, et al. (2014) Acanthamoeba polyphaga mimivirus and other giant viruses: an open field to outstanding discoveries. Virol J 11:120.

40. Reteno DG, et al. (2015) Faustovirus, an asfarvirus-related new lineage of giant viruses infecting amoebae. J Virol.

41. Kenji T, Keizo N, Shigeru I, & Mineo Y (2001) Isolation of a virus infecting the novel shellfish-killing dinoflagellate Heterocapsa circularisquama. Aquat Microb Ecol 23(2):103–111.

42. Caron DA, et al. (2017) Probing the evolution, ecology and physiology of marine protists using transcriptomics. Nat Rev Micro 15(1):6–20.

43. Koch F, Sañudo-Wilhelmy SA, Fisher NS, & Gobler CJ (2013) Effect of vitamins B1 and B12 on bloom dynamics of the harmful brown tide alga, Aureococcus anophagefferens (Pelagophyceae). Limnol Oceanogr 58(5):1761–1774.

44. Guillard RRL (1975) Culture of phytoplankton for feeding marine invertebrates. Culture of Marine Invertebrate Animals, ed Smith WL, Chanley, M.H. (Plenum Press, New York, USA), pp 26–60.

45. Eddy SR (2011) Accelerated profile HMM searches. PLoS computational biology 7(10):e1002195.

46. Keeling PJ, et al. (2014) The marine microbial eukaryote transcriptome sequencing project (MMETSP): illuminating the functional diversity of eukaryotic life in the oceans through transcriptome sequencing. PLoS Biol 12(6):e1001889.

47. Tamura K, Stecher G, Peterson D, Filipski A, & Kumar S (2013) MEGA6: molecular evolutionary genetics analysis version 6.0. Mol Biol Evol 30(12):2725–2729.

48. Guindon S, et al. (2010) New algorithms and methods to estimate maximum-likelihood phylogenies: assessing the performance of PhyML 3.0. Syst Biol 59(3):307–321.

49. Adl SM, et al. (2005) The new higher level classification of eukaryotes with emphasis on the taxonomy of protists. J Euk Microbiol 52(5):399–451.

50. Matsen FA, Kodner RB, & Armbrust VE (2010) pplacer: linear time maximum-likelihood and Bayesian phylogenetic placement of sequences onto a fixed reference tree. BMC Bioinformatics 11(1):1–16.

51. Letunic I & Bork P (2016) Interactive tree of life (iTOL) v3: an online tool for the display and annotation of phylogenetic and other trees. Nucl Acid Res 44(W1):W242–W245.

52. Punta M, et al. (2012) The Pfam protein families database. Nucl Acid Res 40(D1):D290–D301.

53. Marchler-Bauer A, et al. (2013) CDD: conserved domains and protein three-dimensional structure. Nucl Acid Res 41(Database issue):D348–352.

54. Cline MS, et al. (2007) Integration of biological networks and gene expression data using Cytoscape. Nature protocols 2(10):2366–2382.

